# Engineering Substrate Channeling in Assembly-Line Terpene Biosynthesis

**DOI:** 10.1101/2024.03.25.586617

**Authors:** Eliott S. Wenger, Kollin Schultz, Ronen Marmorstein, David W. Christianson

## Abstract

Fusicoccadiene synthase from *P. amygdala* (PaFS) is a bifunctional assembly-line terpene synthase containing a prenyltransferase domain that generates geranylgeranyl diphosphate (GGPP) from dimethylallyl diphosphate (DMAPP) and three equivalents of isopentenyl diphosphate (IPP), and a cyclase domain that converts GGPP into fusicoccadiene, a precursor of the diterpene glycoside Fusicoccin A. The two catalytic domains are linked by a flexible 69-residue polypeptide segment. The prenyltransferase domain mediates oligomerization to form predominantly octamers, and cyclase domains are randomly splayed out around the prenyltransferase core. Previous studies suggest that substrate channeling is operative in catalysis, since most of the GGPP formed by the prenyltransferase remains on the protein for the cyclization reaction. Here, we demonstrate that the flexible linker is not required for substrate channeling, nor must the prenyltransferase and cyclase domains be covalently linked to sustain substrate channeling. Moreover, substrate competition experiments with other diterpene cyclases indicate that the PaFS prenyltransferase and cyclase domains are preferential partners regardless of whether they are covalently linked or not. The cryo-EM structure of engineered “linkerless” construct PaFS_LL_, in which the 69-residue linker is spliced out and replaced with the tripeptide PTQ, reveals that cyclase pairs associate with all four sides of the prenyltransferase octamer. Taken together, these results suggest that optimal substrate channeling is achieved when a cyclase domain associates with the side of the prenyltransferase octamer, regardless of whether the two domains are covalently linked and regardless of whether this interaction is transient or locked in place.

## Introduction

The magnificent chemodiversity of terpenoid natural products is rooted in the simple 5-carbon precursors dimethylallyl diphosphate (DMAPP) and isopentenyl diphosphate (IPP).^1,2^ In the first step of terpene biosynthesis, a prenyltransferase combines DMAPP with one or more equivalents of IPP to form linear isoprenoid products containing 10 carbons (geranyl diphosphate, GPP), 15 carbons, (farnesyl diphosphate, FPP), 20 carbons (geranylgeranyl diphosphate, GGPP), or more in multiples of 5-carbon units.^3,4^ These isoprenoids then undergo multi-step cyclization reactions catalyzed by terpene cyclases to generate complex products typically containing multiple rings and stereocenters.^5–12^ Cyclic terpenes can undergo further oxidation and tailoring reactions to yield bioactive terpenoid products such as the topical anesthetic menthol, the antimalarial drug artemisinin, and the cancer chemotherapeutic drug paclitaxel (Taxol).^13–15^

Assembly-line terpene synthases are bifunctional enzymes containing both prenyltransferase and cyclase domains.^16–20^ These catalytic domains are connected by flexible polypeptide linkers that hinder crystallization, but individual domains can be crystallized for structure determination.^21–23^ For the visualization of full-length enzymes in solution, small-angle X-ray scattering yields low-resolution molecular envelopes showing general domain architecture and quaternary structure.^21,22^ More recently, cryo-electron microscopy (cryo-EM) reveals higher-resolution images of oligomers comprised of unusual combinations of ordered and disordered catalytic domains.^24–26^

Consider the assembly-line terpene synthase, fusicoccadiene synthase from *Phomopsis amygdali* (PaFS), which utilizes DMAPP and 3 equivalents of IPP to generate the tricyclic diterpene fusicoccadiene, which is further derivatized to yield the phytotoxin fusicoccin A (Figure 1A).^16^ Fusicoccin A mediates 14-3-3 protein-protein interactions, and it functions synergistically with interferon-α to induce cancer cell death.^27–29^ PaFS is the first assembly-line terpene synthase to have been discovered,^16^ and it is also the first assembly-line terpene synthase to yield a cryo-EM structure^24^ as well as crystal structures of individual prenyltransferase and cyclase domains.^21^ The individual prenyltransferase domain crystallizes as a hexamer, but cryo-EM and negative-stain EM images of the full-length enzyme reveal a mixture containing approximately 90% octamers and 10% hexamers.^24^ Surprisingly, negative-stain EM images further show that cyclase domains are randomly splayed out around prenyltransferase oligomers, with an average prenyltransferase-cyclase separation of 114 Å.^24^ Even so, 11% of particles contain one or more associated cyclase domains;^26^ this population increases to 45% when protein samples are crosslinked with glutaraldehyde.^24^ Thus, while randomly and dynamically splayed out at any given moment, cyclase domains are capable of transient association with the oligomeric prenyltransferase core as enabled by the 69-residue linker connecting the cyclase to the prenyltransferase.

**Figure 1.**
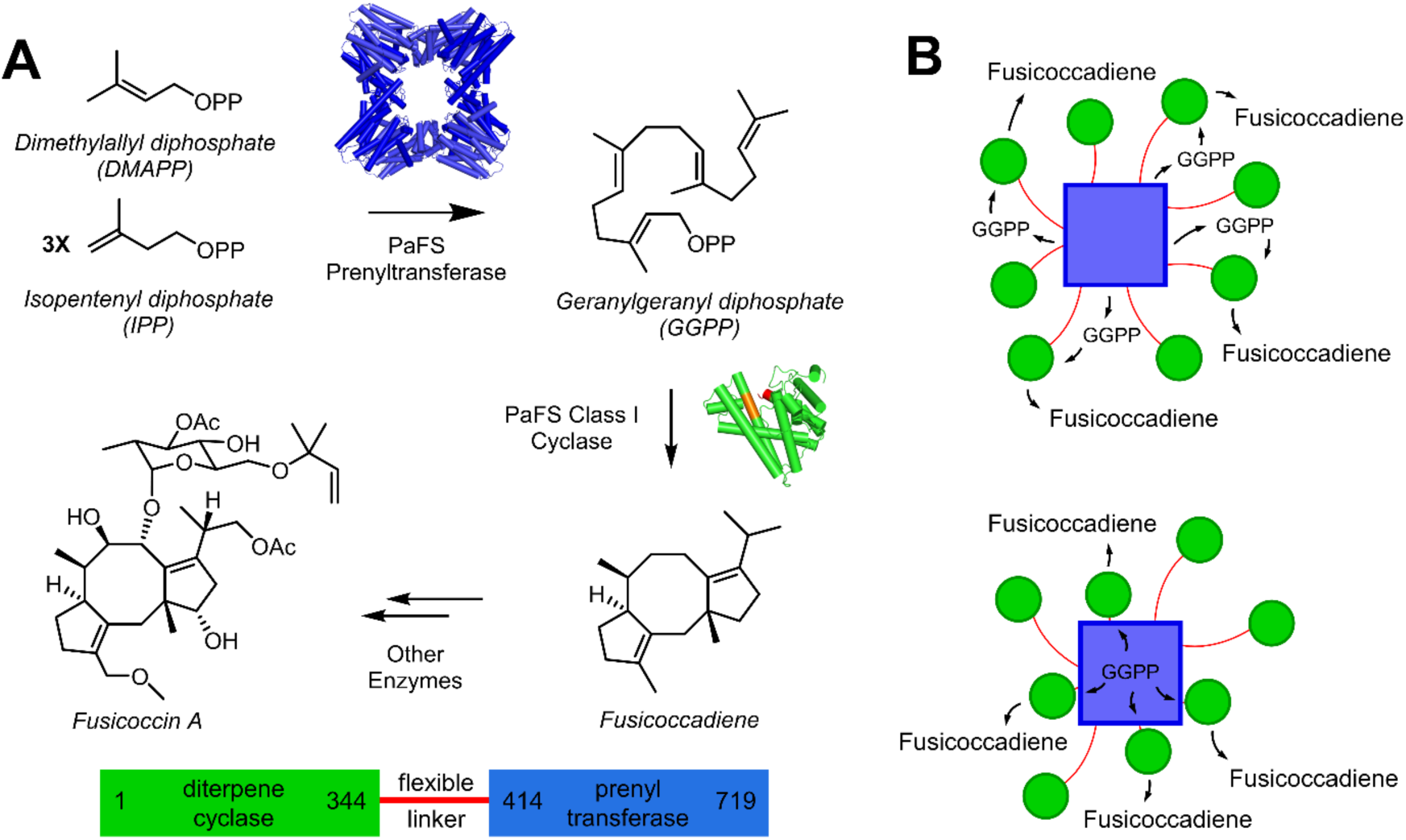
Bifunctional catalysis by PaFS. (A) The octameric prenyltransferase domain catalyzes the processive coupling reaction of one molecule of DMAPP and 3 equivalents of IPP to generate GGPP, which then transits to one of eight cyclase domains to generate fusicoccadiene. Further derivatization of fusiccocadiene yields the phytotoxin fusicoccin A. For reference, the primary structure of PaFS is shown at bottom. (B) Substrate channeling in the PaFS reaction sequence might occur through cluster channeling (top) or by proximity channeling (bottom) enabled by preferential interactions between the octameric prenyltransferase core (blue square) and cyclase domains (green circles).

In view of the dynamic nature of prenyltransferase-cyclase association, it is surprising that most of the GGPP generated in the prenyltransferase core stays on the oligomer for cyclization.^30^ When an equimolar mixture of PaFS and the diterpene cyclase taxadiene synthase is incubated with GGPP, a 5:1 ratio of fusicoccadiene:taxadiene results. However, when incubated with only DMAPP and IPP, such that the only source of GGPP is that generated by the PaFS prenyltransferase core, the fusicoccadiene:taxadiene ratio increases to 48:1 – strikingly, most of the GGPP generated by the PaFS prenyltransferase is directed to the PaFS cyclase despite the competition with taxadiene synthase. Two possible models could account for substrate channeling in this system (Figure 1B). We initially suggested^24^ that cluster channeling^31–33^ could be operative: the probability that a given molecule of GGPP generated by the prenyltransferase encounters a specific cyclase domain is low, but the probability that GGPP encounters any one of eight cyclase domains randomly splayed out around the oligomeric prenyltransferase core is much higher. However, since the cyclase domain is capable of transient association with the side of the prenyltransferase core,^24–26^ additional or alternative modes of substrate channeling might facilitate direct transit of GGPP from the prenyltransferase to the cyclase. Specifically, could GGPP transfer occur when the cyclase domain is transiently docked to the prenyltransferase core?

Here, we probe the molecular basis of GGPP channeling in catalysis by PaFS. Surprisingly, covalent linkage of the prenyltransferase and cyclase domains is not required for GGPP channeling, although covalent linkage does improve channeling efficiency. Moreover, preferential GGPP channeling from the prenyltransferase domain to the cyclase domain of PaFS is observed even when a different cyclase, cyclooctatenol synthase (CotB2),^34,35^ is covalently attached to the PaFS prenyltransferase and allowed to compete with a separate PaFS cyclase construct. These data are consistent with a model in which the PaFS prenyltransferase and cyclase domains transiently associate to enable GGPP channeling, regardless of whether they are covalently linked. We further show that substrate channeling is maintained even when the 69-residue linker of full-length PaFS is spliced out and replaced with the tripeptide PTQ. The cryo-EM structure of this “linkerless” variant (PaFS_LL_) reveals that all eight cyclase domains are locked in place on the sides of the prenyltransferase octamer in positions similar to those observed for associated cyclase domains in 11% of wild-type enzyme particles.^26^ These results suggest that the sides of the prenyltransferase octamer are the preferred locations for cyclase domains to dock – transiently in full-length PaFS, transiently in equimolar mixtures of individual PaFS prenyltransferase and cyclase domains, or permanently in linkerless PaFS – to ensure prenyltransferase-cyclase proximity that supports substrate channeling.

## Results

### Enzyme Constructs and Catalytic Activities

We prepared full-length PaFS, the individual cyclase domain (PaFS_CY_, residues 1-344), and the individual prenyltransferase domain (PaFS_PT_, residues 414-719) as previously described.^21^ We also prepared the prenyltransferase domain of copalyl diphosphate synthase from *Penicillium verruculosum* (PvCPS), a class II assembly-line diterpene synthase, using published procedures.^22^ Additionally, we designed and prepared two newly engineered constructs: “linkerless” PaFS (PaFS_LL_), in which the native 69-residue linker segment is replaced by the tripeptide PTQ; and CotB2-PaFS_PT_, a chimera containing the diterpene cyclase cyclooctatenol synthase (CotB2)^34,35^ followed by the native linker segment and prenyltransferase domain of PaFS. Finally, we prepared the individual class I diterpene cyclases CotB2 and spatadiene synthase (SpS) (Figure S1) according to published procedures.^34,36^

Comparison of steady-state kinetic parameters for all six cyclases reveals generally similar catalytic activities (Figure S2). The three fusicoccadiene synthase constructs (PaFS, PaFS_LL_, and PaFS_CY_) are most efficient, while CotB2 and SpS are slightly less efficient based on *k*_cat_/*K*_M_ (Table S1). The catalytic efficiency of PaFS is similar to that observed in previous studies.^21,24^

### Substrate Competition as an Indicator of Substrate Channeling

We used gas chromatography-mass spectrometry (GC-MS) to study product ratios in GGPP competition experiments between pairs of diterpene cyclases. First, we analyzed substrate competition between full-length PaFS and CotB2 by incubating an equimolar enzyme mixture with either GGPP or DMAPP and IPP (Figure 2A). When incubated with GGPP, the measured product ratio is 6.7:1 fusicoccadiene:cyclooctatenol. However, when incubated with DMAPP and IPP, the product ratio shifts to 52.7:1, reflecting substantial enrichment of fusicoccadiene (Table 1). These results indicate that most of the GGPP generated by the PaFS prenyltransferase remains on the enzyme for cyclization to form fusicoccadiene. In other words, GGPP is channeled from the prenyltransferase to the cyclase, but the channel is not perfect – it is sufficiently leaky to allow some GGPP to escape PaFS for cyclization catalyzed by CotB2.

**Figure 2.**
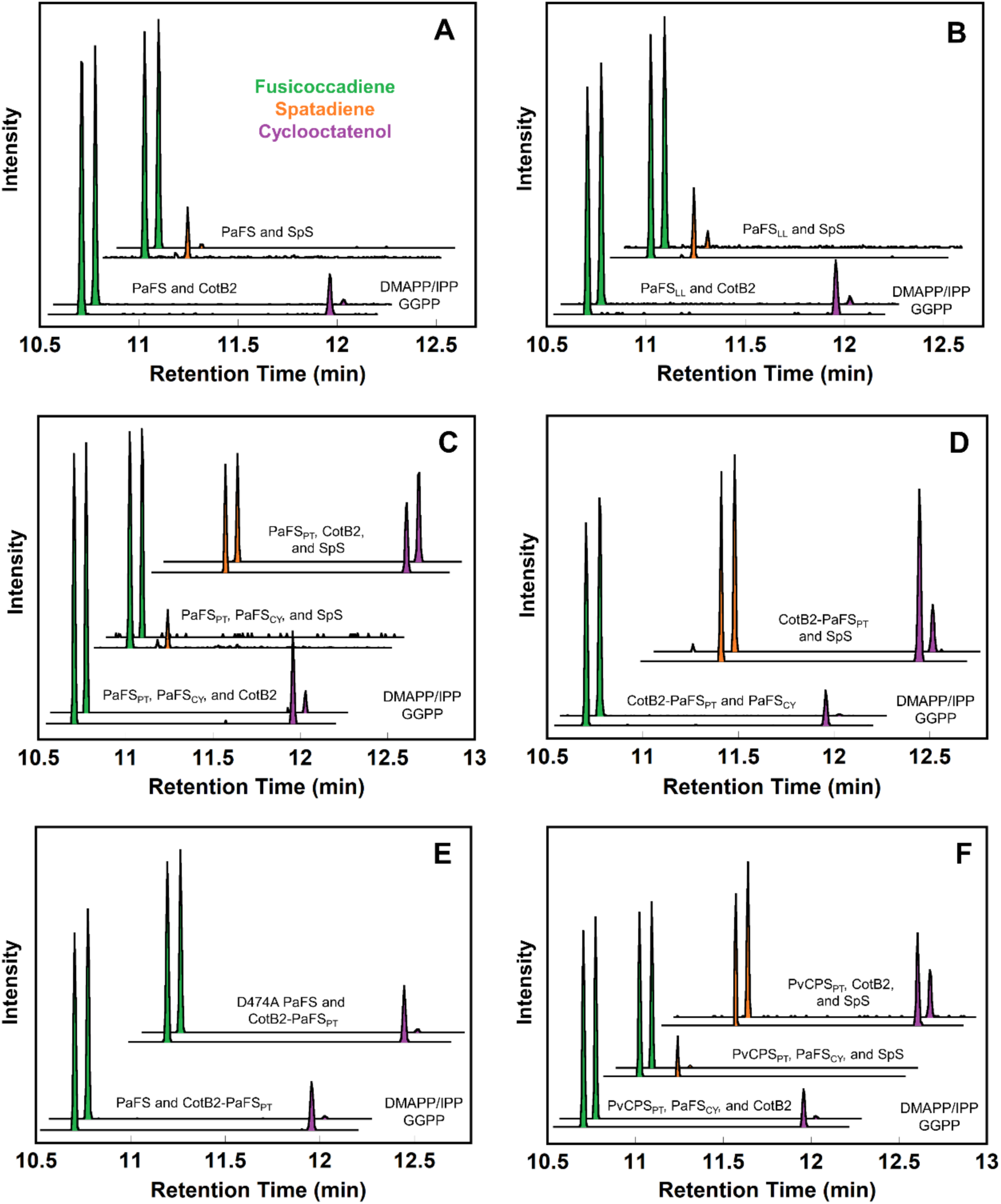
Reaction products generated by enzyme mixtures incubated with GGPP or DMAPP and IPP characterized by GC-MS. (A) Competition between PaFS and CotB2 or SpS. (B) Competition between PaFS_LL_ and CotB2 or SpS. (C) Competition between CotB2-PaFS_PT_ and PaFS_CY_ or SpS. (D) Competition between PaFS_CY_, CotB2, or SpS with PaFS_PT_. (E) Competition between CotB2-PaFS_PT_ and PaFS or PaFS D474A. (F) Competition between PaFS_CY_, CotB2, or SpS with PvCPS_PT_. Each experiment was performed in duplicate; representative chromatograms are plotted. Each set of two traces shows the reaction with GGPP below and the reaction with DMAPP and IPP above. Fusicoccadiene (retention time 10.7 min) peaks are green, spatadiene (retention time 10.9 min) peaks are orange, and cyclooctatenol (retention time 11.9 min) peaks are purple.

**Table 1.** Substrate Competition Assay.

| **Equimolar Enzyme Mixture** | **Products Formed<sup>a</sup>** | **Product Ratio from GGPP** | **Product Ratio from DMAPP and IPP** |
| --- | --- | --- | --- |
| PaFS, CotB2 | FS:CO | $6.7 \pm 0.8 : 1$ | $52.7 \pm 1.6 : 1$ |
| PaFS, SpS | FS:SP | $5.7 \pm 0.4 : 1$ | $74 \pm 9 : 1$ |
| PaFS<sub>LL</sub>, CotB2 | FS:CO | $3.2 \pm 0.4 : 1$ | $27 \pm 5 : 1$ |
| PaFS<sub>LL</sub>, SpS | FS:SP | $4.2 \pm 0.6 : 1$ | $19 \pm 3 : 1$ |
| CotB2-PaFS<sub>PT</sub>, PaFS<sub>CY</sub> | FS:CO | $3.3 \pm 1.7 : 1$ | $83 \pm 5 : 1$ |
| CotB2-PaFS<sub>PT</sub>, SpS | CO:SP | $1.2 \pm 0.1 : 1$ | $0.3 \pm 0.1 : 1$ |
| PaFS<sub>PT</sub>, PaFS<sub>CY</sub>, CotB2 | FS:CO | $3.0 \pm 0.3 : 1$ | $12.9 \pm 1.8 : 1$ |
| PaFS<sub>PT</sub>, PaFS<sub>CY</sub>, SpS | FS:SP | $8.1 \pm 0.7 : 1$ | ND<sup>b</sup> |
| PaFS<sub>PT</sub>, CotB2, SpS | CO:SP | $1.8 \pm 0.7 : 1$ | $1.4 \pm 0.6 : 1$ |
| PaFS, CotB2-PaFS<sub>PT</sub> | FS:CO | $3.4 \pm 0.3 : 1$ | $63.7 \pm 1.7 : 1$ |
| PaFS D474A, CotB2-PaFS<sub>PT</sub> | FS:CO | $2.8 \pm 0.1 : 1$ | $47.9 \pm 1.7 : 1$ |
| PvCPS<sub>PT</sub>, PaFS<sub>CY</sub>, CotB2 | FS:CO | $4.3 \pm 0.3 : 1$ | $42.6 \pm 1.2 : 1$ |
| PvCPS<sub>PT</sub>, PaFS<sub>CY</sub>, SpS | FS:SP | $6.6 \pm 1.4 : 1$ | $68.7 \pm 4.4 : 1$ |
| PvCPS<sub>PT</sub>, CotB2, SpS | CO:SP | $1.2 \pm 0.2 : 1$ | $0.4 \pm 0.1 : 1$ |

<sup>a</sup>FS, fusicoccadiene; CO, cyclooctatenol; SP, spatadiene. <sup>b</sup>ND, spatadiene not detected.

To further confirm that the enrichment of fusicoccadiene generation is not dependent on the diterpene cyclase that competes for GGPP, we repeated the competition experiment with an alternative cyclase, SpS (Figure 2A). When an equimolar mixture of PaFS and SpS is incubated with GGPP, the measured product ratio is 5.7:1 fusicoccadiene:spatadiene; when incubated with DMAPP and IPP, the product ratio shifts to 73:1, again reflecting substantial enrichment of fusicoccadiene (Table 1). Here, too, most of the GGPP generated by the PaFS prenyltransferase stays on the enzyme for cyclization to form fusicoccadiene, indicative of substrate channeling.

We next measured product generation by the “linkerless” construct PaFS_LL_, which retains normal fusicoccadiene synthase activity (Table S1, Figure S3). When PaFS_LL_ is subjected to competition with CotB2 or SpS, enrichment of fusicoccadiene production is observed when the equimolar enzyme mixture is incubated with DMAPP and IPP, although the magnitude of enrichment is somewhat diminished compared with full-length PaFS (Table 1, Figure 2B). Thus, substrate channeling is preserved in PaFS_LL_, although optimal channeling requires the flexible 69-residue linker between the prenyltransferase and the cyclase domains.

Must the prenyltransferase and cyclase of PaFS be covalently linked to support substrate channeling? An equimolar mixture of PaFS_PT_, PaFS_CY_, and CotB2 incubated with GGPP yields a product ratio of 3:1 fusicoccadiene:cyclooctatenol (Table 1, Figure 2C). However, when incubated with DMAPP and IPP, the ratio shifts to 13:1 fusicoccadiene:cyclooctatenol. An even larger shift is observed when studying an equimolar mixture of PaFS_PT_, PaFS_CY_, and SpS: the product ratio measured after incubation with GGPP is 8:1 fusicoccadiene:spatadiene, but when incubated with DMAPP and IPP, negligible spatadiene is detected while a similar amount of fusicoccadiene is observed. Therefore, the PaFS prenyltransferase and cyclase domains need not be covalently linked to achieve substrate channeling, but the magnitude of channeling is diminished compared to that observed for wild-type PaFS and PaFS_LL_.

To confirm that substrate channeling is specific to the pairing of PaFS prenyltransferase and cyclase domains, we studied an equimolar mixture of PaFS_PT_, CotB2, and SpS. When incubated with GGPP or DMAPP and IPP, the measured product ratios are essentially invariant (Table 1, Figure 2C). We also studied the chimeric bifunctional enzyme CotB2-PaFS_PT_ in competition with individual PaFS_CY_ or SpS cyclases (Figure 2D). When incubated with GGPP, the measured product ratio is 1.9:1 fusicoccadiene:cyclooctatenol, and when incubated with DMAPP and IPP the product ratio shifts substantially to 75:1 fusicoccadiene:cyclooctatenol (Table 1). Surprisingly, fusicoccadiene synthesis from the GGPP generated by the PaFS prenyltransferase domain is preferred even when the individual PaFS cyclase domain is an intermolecular competitor for the GGPP generated by the prenyltransferase domain of CotB2-PaFS_PT_. Consistent with this observation, when an equimolar mixture of CotB2-PaFS_PT_ and SpS is incubated with GGPP or DMAPP and IPP, the measured product ratio does not substantially shift (Table 1, Figure 2D). Thus, the PaFS cyclase is the preferred partner of the PaFS prenyltransferase regardless of whether or not these catalytic domains are covalently linked. Absent PaFS_CY_, there is no substrate channeling from PaFS_PT_ to non-native cyclases.

To further interrogate the preferential interaction between the prenyltransferase and cyclase domains of PaFS, we assayed an equimolar mixture of two different bifunctional constructs, wild-type PaFS and CotB2-PaFS_PT_ (Figure 2E). Analysis of 1 μM protein samples using size exclusion chromatography indicates heterooligomerization, such that each octamer generally consists of 4 wild-type PaFS chains and 4 CotB2-PaFS_PT_ chains (Figure S4). When incubated with GGPP, the measured product ratio is 3.4:1 fusicoccadiene:cyclooctatenol, not too different from the ratio measured when equimolar PaFS and CotB2 are incubated with GGPP (6.7:1). When the heterooctamer is incubated with DMAPP and IPP, the product ratio shifts to 64:1 fusicoccadiene:cyclooctatenol, slightly better enrichment than that of 53:1 measured when measured when equimolar PaFS and CotB2 are incubated with these 5-carbon precursors.

Similar observations result when studying an equimolar mixture of CotB2-PaFS_PT_ and a full-length PaFS variant containing an inactivated prenyltransferase domain (PaFS D474A) (Figure 2E). These constructs are similarly presumed to form heterooligomers consisting of 4 PaFS D474A chains and 4 CotB2-PaFS_PT_ chains When incubated with GGPP, the measured product ratio is 3:1 fusicoccadiene:cyclooctatenol, but when incubated with DMAPP and IPP, the product ratio shifts to 48:1. Thus, optimal substrate channeling occurs between the cognate prenyltransferase and cyclase domains of PaFS, even if the PaFS cyclase domain is not contained in the same subunit as the catalytically active PaFS prenyltransferase domain.

With a preferential functional interaction between the prenyltransferase and cyclase domains of PaFS thus established, we tested whether utilizing a different prenyltransferase might disrupt this interaction (Figure 2F). When an equimolar mixture of PaFS_CY_, CotB2, and the hexameric prenyltransferase domain of PvCPS (PvCPS_PT_) is incubated with GGPP, the measured product ratio is 4:1 fusicoccadiene:cyclooctatenol. When incubated with DMAPP and IPP, the ratio shifts to 43:1 fusicoccadiene:cyclooctatenol (Table 1). Surprisingly, the preferential interaction established between the prenyltransferase and cyclase domains of PaFS is conserved between the prenyltransferase domain of PvCPS and the cyclase domain of PaFS. Fusicoccadiene enrichment was also observed when this experiment was repeated with an alternative cyclase competitor, SpS. Here, too, fusicoccadiene enrichment is observed (Table 1, Figure 2F). When the two non-assembly-line cyclases CotB2 and SpS compete for GGPP generated by PvCPS_CT_ in an equimolar enzyme mixture, only a negligible change in the product ratio is observed (Table 1, Figure 2F). Since PvCPS is a class II assembly-line terpene synthase, it is possible that the structural features facilitating transient prenyltransferase-cyclase association between PaFS_PT_ and PaFS_CY_ are conserved in PvCPS_PT_. This would account for the preferential channeling of GGPP to PaFS_CY_, as well as the lack of observed channeling in cyclase mixtures lacking PaFS_CY_.

### Cryo-EM Structure of PaFS_LL_

To provide a framework for understanding substrate channeling in an engineered “linkerless” assembly-line terpene synthase, we determined the cryo-EM structure of PaFS_LL_. Micrographs of grids prepared with 0.8 mg/mL protein samples yield two-dimensional (2D) classes with predominantly octameric quaternary structure, although hexamers are also observed (Figure S5). Facile interchange of oligomeric species might thus be expected in solution, as also suggested by the observation of multiple oligomeric species using mass photometry at 10^3^-fold lower protein concentrations (Figure S6).

In contrast with the cryo-EM structure of wild-type PaFS,^24,26^ the cyclase domains of PaFS_LL_ are readily visualized in 2D classes. The best 2D classes of the PaFS_LL_ octamer show eight well-ordered cyclase domains associated with the sides of the prenyltransferase core – one cyclase pair on each side of the octamer, such that cyclase pairs appear to be flat or nearly flat in the plane of the prenyltransferase core (Figure S5A). However, other 2D classes show that cyclase pairs can tilt up to ∼60° or more (Figure S5B).

After inspection of 2D class averages, we proceeded with the three-dimensional (3D) structure determination of octameric PaFS_LL_. Data collection and refinement statistics are listed in Table S2 and the workflow of the structure determination is summarized in Figure S7. Using 220,663 octamer particles, we generated a 3D reconstruction of the octameric prenyltransferase at 3.56 Å resolution, which improved slightly to 3.53 Å resolution when cyclase domains were excluded from refinement using particle subtraction (Figure 3, Figure S8A).

**Figure 3.**
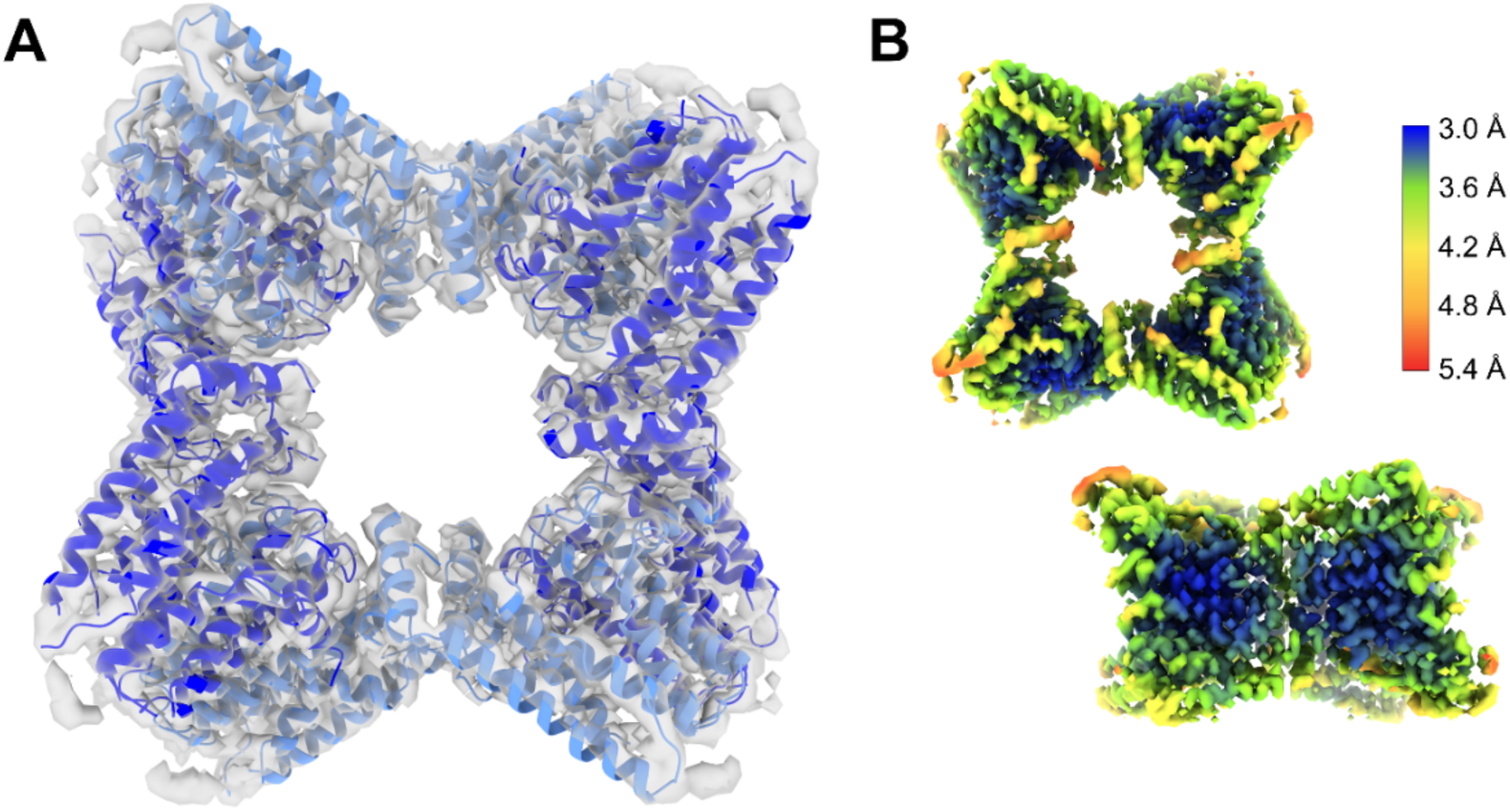
Cryo-EM structure of the PaFS_LL_ octameric prenyltransferase core. (A) Map of the PaFS_LL_ prenyltransferase octamer with *C*2 symmetry fit with atomic coordinates. (B) Local resolution estimation for top and side views of the octamer.

The quaternary structure of the prenyltransferase octameric core is essentially identical to that of wild-type PaFS. At first glance, the overall symmetry of the octamer appears to be *D*4, but local structural differences result in overall *C*2 symmetry. The tertiary structure of the prenyltransferase domain is essentially identical to that observed in wild-type PaFS, with an overall root-mean-square (rms) deviation of 0.88 Å for 214 Cα atoms between monomer A of wild-type PaFS and monomer A of PaFS_LL_; relatively minor differences in quaternary structure are observed (Figure S9). Thus, splicing out the linker does not perturb the structure of the prenyltransferase core. Prenyltransferase active sites open into the central pore of the octamer.

To better understand the positioning of cyclase pairs around the prenyltransferase octamer, we subjected the pool of octamer particles to simple 3D classification or heterogeneous refinement, the latter of which allows for the rotation of particles during classification. Strikingly, a class with all four cyclase pairs in the flat orientation did not emerge from either approach. Instead, classes contained mixtures of cyclase orientations in different combinations. Therefore, we turned to 3D variability analysis to solve for 3 modes of continuous motion, to generate the best representation of the entire particle set. Along the component that described the majority of the motion in the dataset, we generated 20 maps illustrating a transition from a mostly compact conformation with tilted cyclase pairs to a more expanded conformation with 3 out of 4 cyclase pairs tending toward a flat orientation (Movie 1, available online). Snapshots from this movie are illustrated in Figure 4.

**Figure 4.**
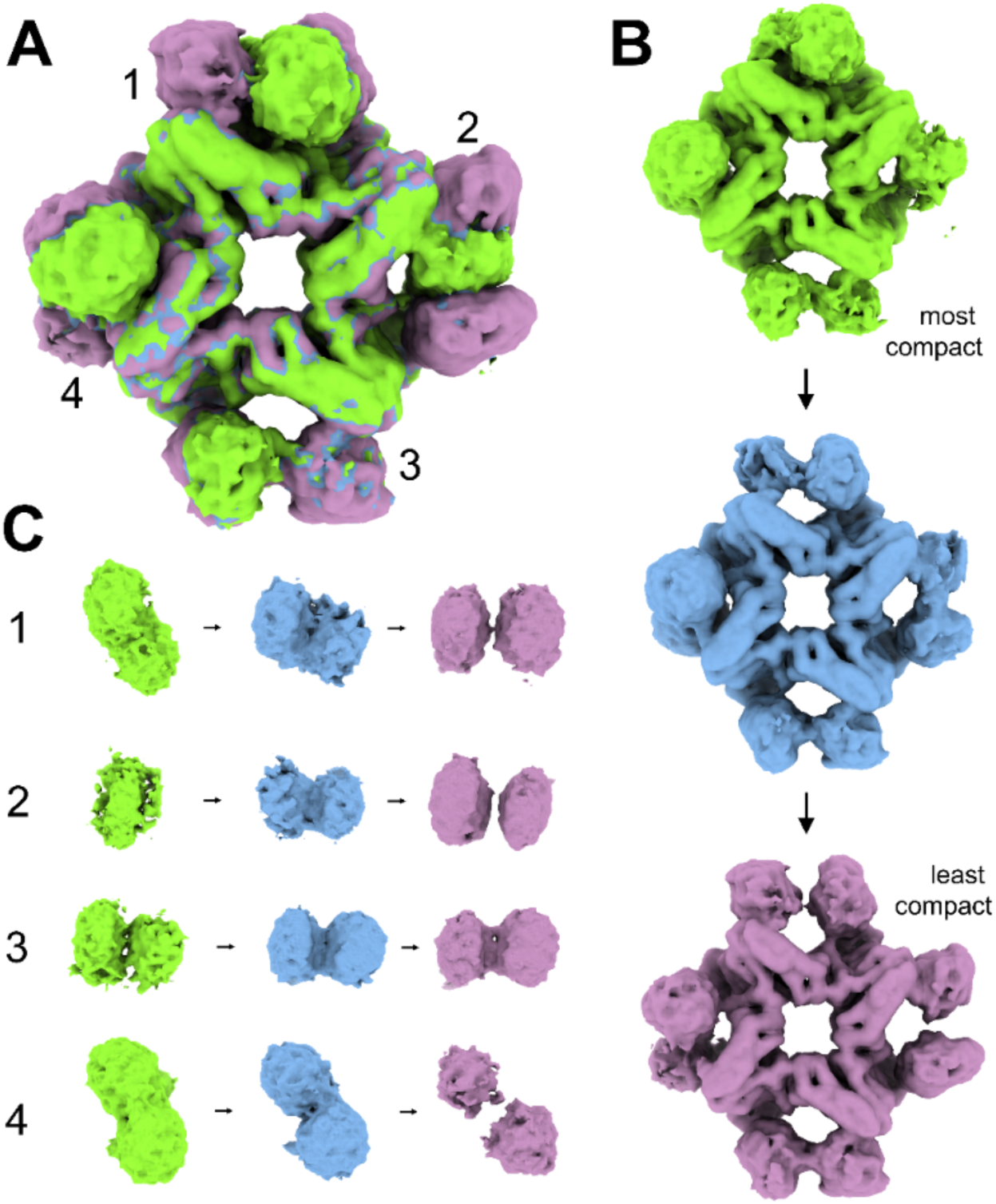
3D conformational variability analysis. Three maps of the 20 total appearing in Movie 1 summarize the conformational transition of PaFS_LL_ from the most compact state (green), through an intermediate state (blue), to the most expanded state (purple). (A) Overlay of the three maps. (B) Individual top-down views of each map. (C) Side views of cyclase pairs 1–4. Generally, cyclase pairs appear to shift in concert from compact/tilted to expanded/flat conformations. However, closer inspection reveals that each cyclase pair exhibits slightly different behavior. Pair 1 shifts from tilted to flat, while pair 2 shifts from a single domain in the plane of the octamer (possibly representing a mixture of folded and denatured states) to a flat pair with clear separation. Pair 3 is flat and shows the least movement, while pair 4 is tilted through the whole series but grows flatter and gains clear separation into two separate domains.

Finally, seeking to visualize cyclase domains at the highest resolution possible, we subjected all octamer particles to 3D classification with a focus mask on the cyclase domains. Because the program aligns one well-ordered cyclase pair per octamer, density for the remaining three cyclase pairs in each class becomes correspondingly less well-observed, as a mixture of various orientations are averaged together. The best class depicted a flat cyclase pair, while the densities of tilted cyclase pairs were not as well defined. The remaining three classes depict cyclase pairs at intermediate tilt angles or with partial or nearly complete denaturation of one or both of the cyclase domains in the pair.

Ultimately, we generated a 4.79 Å-resolution map of an octameric prenyltransferase core with two interacting cyclase domains in the flat orientation (Figure 5A-B; workflow summarized in Figure S7 and FSC plot recorded in Figure S8). We modeled six of the prenyltransferase chains as free prenyltransferase domains and modeled the entire sequence of full-length PaFS_LL_ into the remaining di-domain densities. Intriguingly, the cyclase domains are not symmetrically associated with the octameric core. One is more well-ordered than the other (denoted Cyclase 1), with clear density appearing for the engineered PTQ linker segment (connects to red helix at bottom) (Figure 5C). The other (Cyclase 2) appears to associate in a slightly different manner, and while the density is sufficient to clearly define an orientation for the docked chain, no density is observable at the C-terminus where the domain connects to the octameric core.

**Figure 5.**
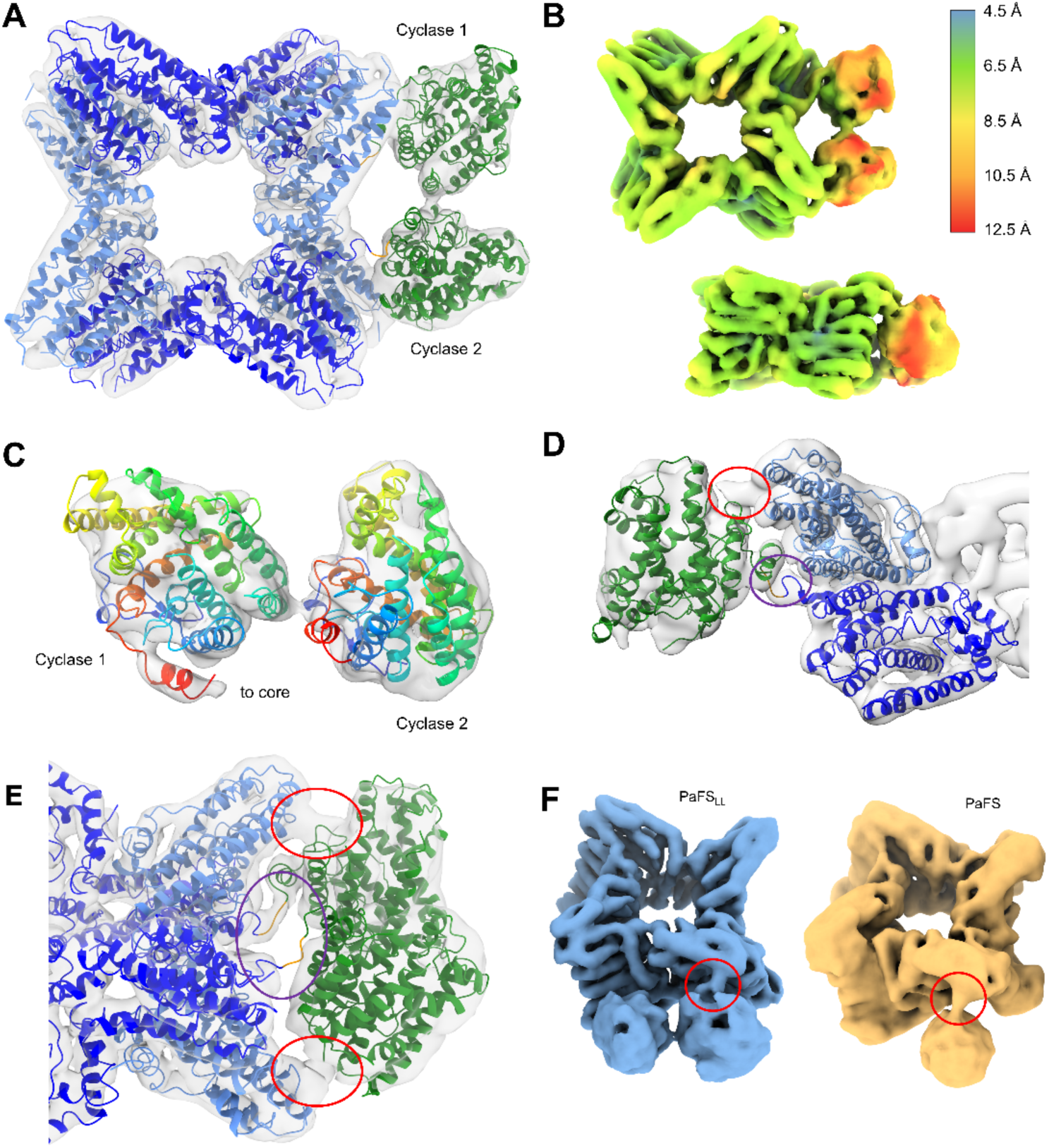
Structure of the PaFS_LL_ octamer with an associated cyclase pair. (A) The model was generated by docking the crystal structure of PaFS_CY_ (green; PDB 5ERM^21^) into density for each cyclase domain, then building the linker segment (orange) to connect with the prenyltransferase domain (blue). (B) Local resolution estimation for the octameric core and cyclase pair; top and side views. (C) Cyclase domains adopt slightly different orientations relative to the core; density for one orientation (cyclase 1) was slightly better defined than that for the other (cyclase 2). The chains are colored blue (N-terminus) to red (C-terminus; connection to PaFS_PT_) to facilitate visual comparison of their orientation. (D) Close-up of the linker region of Cyclase 1, with the linker residues colored orange, revealing that prenyltransferase-cyclase association is achieved through domain-swapping: Cyclase 1 (green chain) forms an interaction typified by continuous density (red circle) with a prenyltransferase domain (light blue chain) other than the one it is covalently linked to (dark blue chain, purple circle). (E) Even though there is only visible density for one of the two modeled linker regions (purple circle), both cyclase domains interact with prenyltransferase domains at a symmetrically-related site at the corner of the octameric assembly (red circles). (F) This site is the same as the one identified for inter-domain interactions in wild-type PaFS (EMD-28076), which bears the full-length linker region (red circles).^26^

The model of the prenyltransferase octamer with two associated cyclase domains reveals that prenyltransferase-cyclase association occurs through domain swapping. In other words, while cyclase domain A (green) is covalently linked to prenyltransferase domain A (dark blue), cyclase domain A interacts mainly with prenyltransferase domain B (light blue) (Figure 5D). Continuous density is also observed between cyclase domain A and prenyltransferase domain B at an additional site corresponding to that previously identified in the cryo-EM structure of full-length PaFS (Figure 5E-F). Thus, truncation of the linker segment from 69 residues to 3 residues serendipitously fixes the cyclase domain in the same location as that observed in the full-length enzyme that facilitates GGPP channeling from the prenyltranferase to the cyclase.

## Discussion

While enzymes and metabolites can freely diffuse to a homogenous distribution within a given cellular compartment in the absence of active transport, sequential enzymes in a metabolic pathway, or multiple domains of a polyfunctional enzyme catalyzing sequential reactions, can sometimes associate to form molecular channels that confine and guide metabolites from one enzyme to another, thereby improving pathway efficiency.^37–39^ The presence of such a molecular channel is readily discerned from inspection of an experimentally determined protein structure; however, does channeling always require a channel?

The answer to this question is “no”, since simple proximity can improve pathway efficiency in tandem enzyme-catalyzed reactions. Calculations based on rates of diffusion suggest that enzyme active sites within 10 Å of each other will enable proximity channeling to improve pathway efficiency, i.e., such that the intermediate product released by the first enzyme is not entirely lost to bulk solution but can diffuse directly into the active site of the second enzyme.^40^ However, experiments show that enhanced pathway efficiency can be achieved when enzyme active sites are separated by even greater distances, ∼50 Å or greater.^41,42^ Enzyme partners need not be tightly bound to each other to achieve channeling and improve pathway efficiency – weakly-associated enzyme clusters (metabolons) enhance efficiency in a variety of pathways,^43^ including the tricarboxylic acid cycle in beef heart,^44^ purine nucleotide biosynthesis in humans,^45^ and glucosinolate biosynthesis in plants.^46^

If multiple copies of both enzymes catalyzing a tandem reaction sequence cluster together, the probability may be higher that the intermediate generated by the first enzyme will encounter any one of the second enzymes before escaping to bulk solution.^31^ Thus, channeling does not always require a channel, nor does it require a stable, static multi-enzyme complex – transiently-formed, dynamic multi-enzyme complexes or aggregates can also support channeling.^32^

Here, we provide key insight regarding substrate channeling in a bifunctional assembly-line terpene synthase. Substrate competition experiments interpreted in view of the cryo-EM structures of the wild-type PaFS octamer^24,26^ and the PaFS_LL_ octamer (Figure 5) are consistent with a model for catalysis in which GGPP transit occurs when the cyclase domain is transiently docked at the side of the prenyltransferase octamer (Figure 1B). Our reasoning can be summarized in three major points and illustrated in cartoon form in Figure 6.

**Figure 6.**
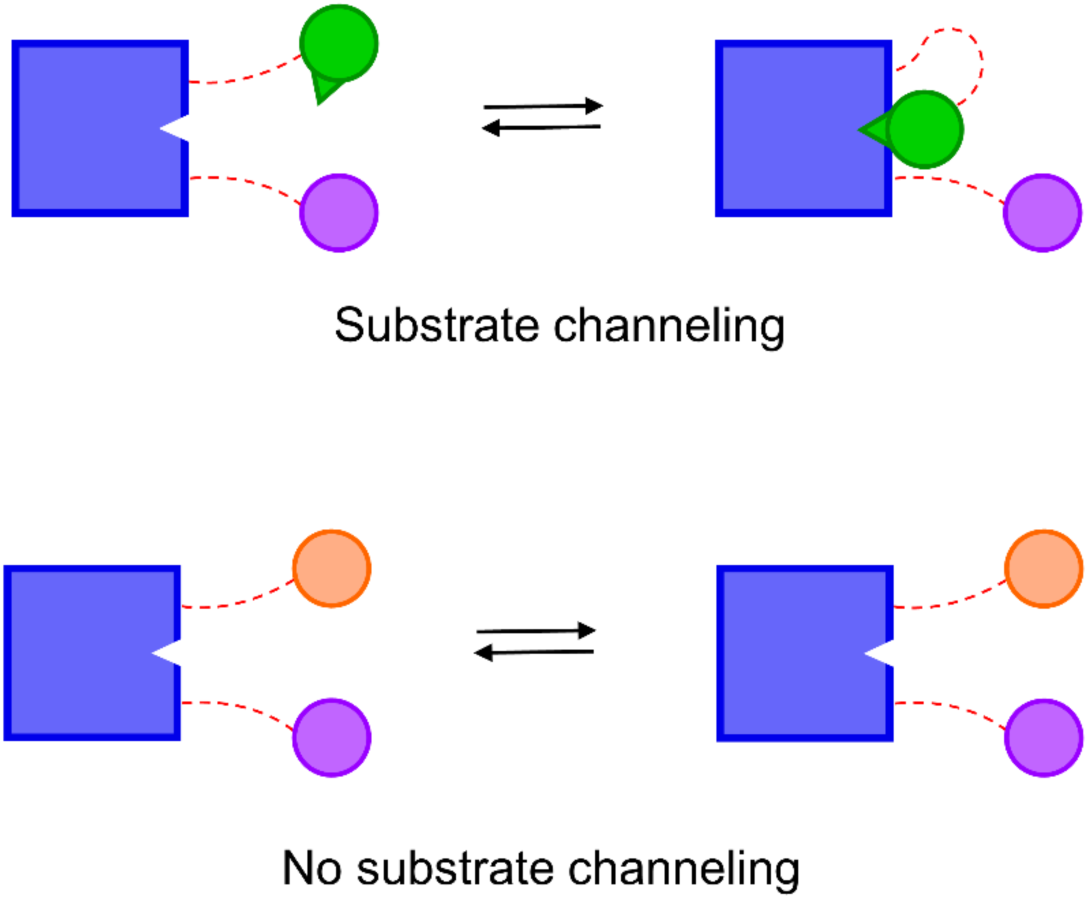
Cartoon summarizing results of substrate channeling experiments. Our experiments suggest that a preferred interaction between the PaFS prenyltransferase octamer (blue square) and a cyclase domain (green circle) facilitates substrate channeling, even when the prenyltransferase and cyclase domains are not covalently linked (the flexible linker is represented by a dashed line to denote that it can be present or absent). Non-native cyclases CotB2 (purple circle) and SpS (orange circle) do not engage in substrate channeling and hence are unable to make a preferred interaction with the PaFS prenyltransferase octamer, regardless of whether they are covalently linked or not.

First, substrate competition experiments universally reveal substantial enrichment of fusicoccadiene generation by full-length PaFS when DMAPP and IPP instead of GGPP are provided to an equimolar mixture of PaFS and a diterpene cyclase competitor – taxadiene synthase,^30^ cyclooctatenol synthase, or spatadiene synthase (Table 1). When DMAPP and IPP are utilized as substrates, the only source of GGPP is that generated by the prenyltransferase domain of PaFS; since fusicoccadiene enrichment is observed under these circumstances, most of the GGPP generated by the PaFS prenyltransferase must stay on the enzyme for cyclization to fusicoccadiene.

Second, substantial enrichment of fusicoccadiene generation with DMAPP and IPP versus GGPP is observed even when the PaFS prenyltransferase and cyclase domains are not covalently linked, although the enrichment is less that that observed when the domains are connected. Even if a different cyclase is attached to the PaFS prenyltransferase and linker, as in CotB2-PaFS_PT_, fusicoccadiene enrichment is observed when this construct competes with PaFS_CY_ for GGPP. If a simple proximity effect were operative, enrichment of cyclooctatenol would be expected. Given that (1) cyclase domains associate predominantly with the sides of the prenyltransferase octamer in full-length PaFS,^24,26^ (2) cyclase domains are locked in similar locations on the sides of the prenyltransferase octamer of PaFS_LL_, and (3) fusicoccadiene enrichment is comparable for full-length PaFS and PaFS_LL_ in substrate competition experiments, the side of the PaFS prenyltransferase octamer is the preferred binding site of the PaFS cyclase domain regardless of whether they are covalently linked or not. Cyclase association with the side of the prenyltransferase octamer supports channeling regardless of whether cyclase-prenyltransferase association is transient, as for full-length PaFS, or static, as for PaFS_LL_.

Third, substantial fusicoccadiene enrichment is observed in an equimolar mixture of CotB2-PaFS_PT_ and PaFS-D474A, a mutation that inactivates the prenyltransferase domain but does not hinder prenyltransferase-mediated oligomerization. Preferential association of the four PaFS cyclase domains from PaFS-D474A with the sides of the (CotB2-PaFS_PT_)_4_(PaFS-D474A)_4_ heterooctamer would still support GGPP channeling from the four active prenyltransferase domains of CotB2-PaFS_PT_. These results also rule out a channeling model in which GGPP is funneled directly from a specific prenyltransferase domain to a specific cyclase domain.

What is the functional importance of the linker, then, that connects the prenyltransferase and the cyclase in PaFS and other assembly-line terpene synthases? Since the channeling efficiencies of full-length PaFS and PaFS_LL_ are roughly the same, the linker is clearly not required for channeling. Nor does the linker substantially influence catalysis by the cyclase domain, since the catalytic efficiency (*k*_cat_/*K*_M_) of PaFS_CY_ is only slightly lower than that of full-length PaFS when incubated with GGPP (Table S1). While the linker can accommodate an alternative cyclase, as exemplified by CotB2-PaFS_PT_, the preferred cyclase partner to receive PaFS_PT_-generated GGPP is always the native cyclase PaFS_CY_. Perhaps the function of the linker is simply to enable the co-evolution of prenyltransferase and cyclase domains in a single polypeptide chain as preferred partners, and thence to ensure that these partners remain close. Alternatively, or additionally, the function of the linker requires stoichiometric expression so as to enable efficient biosynthetic flux at low copy number, avoiding the generation of excess prenyltransferase or cyclase, a possible strategy underlying the covalent linkage of catalytic domains in fatty acid synthase.^47^

In summary, this study illuminates a fascinating example of substrate channeling in complex hydrocarbon biosynthesis. Our model for channeling implicates two catalytic domains that evolved together to sustain the ability for substrate transfer through transient association. In this sense, the nature of the linker is essentially irrelevant – as long as two independent catalytic domains are covalently linked in the same polypeptide chain, they function together and their co-evolution is inextricably linked. At present, it is not possible to “short circuit” the preferential association of PaFS_PT_ and PaFS_CY_ with other cyclases such as CotB2 or SpS to displace PaFS_CY_ as the preferred recipient of GGPP from PaFS_PT_. Even if PaFS_CY_ is utilized as an intermolecular competitor to the potential intramolecular transit of GGPP from PaFS_PT_ to CotB2 in the CotB2-PaFS_PT_ chimera, the preferential generation of fusicoccadiene is sustained, implying preferential association of PaFS_CY_ with PaFS_PT_. Continued investigation of the specific residues that mediate this special association may allow for the engineering of preferred prenyltransferase-cyclase pairs for the efficient biosynthesis of myriad terpene scaffolds important for medicine and biotechnology.

## Acknowledgments

We thank Drs. Sudheer Molugu and Stefan Steimle of the Beckman Center for Cryo-Electron Microscopy (RRID: SCR_022375), University of Pennsylvania, for technical assistance and advice with sample preparation and data collection. Additionally, we thank the NIH for grant R01 GM56838 to D.W.C. and R35 GM118090 to R.M. in support of this research. K.S. was supported by NIH Chemistry-Biology Interface Training Grant T32 GM133398.

## Author Contributions

**Eliott S. Wenger:** Methodology, Investigation, Formal analysis, Validation, Writing - original draft, Writing - review & editing, Visualization.

**Kollin Schultz:** Methodology, Validation, Formal analysis, Writing - review & editing.

**Ronen Marmorstein:** Formal analysis, Resources, Writing - review & editing, Funding acquisition.

**David W. Christianson:** Conceptualization, Formal analysis, Resources, Writing - review & editing, Project administration, Funding acquisition.

## Data Availability

Atomic coordinates of the PaFS_LL_ prenyltransferase octamer (*C*2 symmetry) have been deposited in the Protein Data Bank (PDB, www.rcsb.org) with accession code 9B3T, and the cryo-EM map of this structure has been deposited in the Electron Microscopy Data Bank (EMDB) with accession code EMD-44155. The asymmetric (*C*1) cryo-EM map showing two cyclases associated with the octameric prenyltransferase core has also been deposited in the EMDB with accession code EMD-44098. The previously-determined^21^ crystal structure of the individual PaFS cyclase domain was used in this study and can be accessed in the PDB with accession code 5ERM. The previously determined^24^ cryo-EM structure of the PaFS octameric prenyltransferase core (PDB 8EAX, EMD-27989) and the map with one associated cyclase domain (EMD-28076) were also used in this study. All other relevant data are available from the corresponding author upon request. Source data are provided with this paper.

## Methods

### Protein Expression and Purification

All proteins described herein were prepared in similar fashion. Plasmids containing the gene of interest fused to an N-terminal His6 tag in the pet28A vector were obtained from Genscript and transformed into BL21-DE3 competent cells (New England Biolabs). After growth overnight at 37°C on an agar plate, a single colony was used to inoculate a starter culture supplemented with 50 µg/mL kanamycin. The starter culture was grown overnight at 37°C with shaking and then used to inoculate 6 x 1-L LB media containing 50 µg/mL kanamycin. Once the optical density at 600 nm reached 1.0, flasks were incubated on ice for 45 min and then induced with 0.5 mM isopropyl β-D-1-thiogalactopyranoside (IPTG). After shaking at 16°C for 18 h, cells were pelleted by centrifugation and resuspended in lysis buffer [50 mM 4-(2-hydroxyethyl)-1-piperazineethanesulfonic acid (HEPES) (pH 7.5), 200 mM NaCl, 2 mM tris(2-carboxyethyl)-phosphine hydrochloride (TCEP), and 10% glycerol] and stirred for two hours (all purification steps were performed at 4°C). Cells were then lysed by sonication (1 s-on/3 s-off at 30% power for 40 min) and clarified by centrifugation.

Supernatant was loaded onto a Ni-NTA column (Cytiva) pre-equilibrated with 5 column-volumes of lysis buffer; protein was eluted using elution buffer (lysis buffer with 300 mM imidazole). Elution fractions containing the most protein were combined and loaded directly onto a size-exclusion column (GE Healthcare) pre-equilibrated with lysis buffer without prior concentration or dialysis of the protein. Fractions containing the most protein were combined and diluted to 100 mL total volume using zero-salt lysis buffer (lysis buffer containing no NaCl) and loaded onto an anion-exchange column (Cytiva) pre-washed with 5 column-volumes of zero-salt buffer. Protein was eluted with a 100-mL gradient between the zero-salt buffer and high-salt buffer (lysis buffer containing 1 M NaCl). Protein fractions were flash-cooled in liquid nitrogen and stored in a -80 °C freezer without being pooled to preserve the high protein concentrations in the peak fractions.

### Steady-State Kinetics

Enzyme activity was measured using the EnzChek^TM^ Pyrophosphate Detection Assay Kit (ThermoFisher Scientific) as previously described for PaFS.^24^ Cyclization assays were performed in duplicate on a 100-µL scale with the protein concentration at 500 nM in lysis buffer. MgCl_2_, MESG, PNP, and IPP were added to final concentrations of 2.5 mM, 0.2 mM, 0.1 U, and 0.003 U, respectively. Each reaction was individually assembled in a cuvette to a final volume of 90 µL, containing the appropriate concentration of GGPP (ranging from 2.5 to 50 µM) added from the working dilution (a 5 mM stock was made by reconstitution of solid in 70% methanol, 30% 10 mM NH_4_HCO_3_ in water at 5 mM and the 0.5 mM working dilution was made by dilution to 0.5 mM in lysis buffer). The reaction was then initiated by the addition of 10 µL of enzyme from a 5 µM stock. The cuvette was then placed into a Cary 60 UV/vis spectrometer and the absorbance at 360 nm was measured every 2 s for 2 mins. In total, two replicates of nine different substrate concentrations were measured for each of the six enzymes.

We determined the initial velocity of each reaction by fitting the linear portion of each trace (40–80 s) using linear regression in Prism software. Slopes were converted to rates (rise in AU/s^-1^ to nM phosphate/s^-1^) using the conversion factor (0.0258) determined from a control experiment where pyrophosphate was added directly to the assay. The rates were then plotted over GGPP concentration and analyzed using Prism software. Data were fit to the Michaelis-

Menten equation:

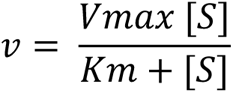

Catalytic parameters are recorded in Table S1.

### Substrate Competition Assay

In a typical substrate competition experiment, both proteins were diluted to 3.5 µM in freshly-prepared lysis buffer with a total volume of 225 µL in a 2-mL glass vial with no insert. MgCl_2_ was added to a final concentration of 2.5 mM. To initiate the reaction, 22.5 µL of substrate was added – either GGPP from a 5 mM stock, or 12.5 µL of DMAPP from a 10 mM stock and 12.5 µL of IPP from a 30 mM stock. The solution was quickly overlaid with 250 µL of hexanes and the vial was capped. All substrates were resuspended in a mixture of 70% methanol, 30% 10 mM NH_4_HCO_3_ in water, and adding substrate in the specified amounts from the specified stock concentrations ensured that each sample contained the same final volume of methanol regardless of which substrates were added. The vial was incubated for 4 h at room temperature, and then vortexed for 15 s in 5-s increments. Because the activity of PaFS_PT_ purified as an individual domain is lower than that measured using full-length PaFS, competition assays utilizing PaFS_PT_ were incubated for 14 h instead of 4 h. The organic layer (100 µL) was then removed using a micro-pipette and transferred to a fresh GC vial with an insert.

Gas chromatography-mass spectrometry (GC-MS) was used for the identification and quantification of diterpene cyclization products. Samples were then run on an Agilent 8890 GC system coupled to a 5977C GC/MSD mass spectrometer with a J&W HP-5MS GC column capillary column in EI+ mode. The following temperature program was used: hold at 60°C for 2 min, ramp to 320°C at 10°C/min, and hold at 320°C for 2 min. A solvent delay of 4 min was used. Fusicoccadiene eluted at 10.7 min, spatadiene at 10.9 min, and cyclooctatenol at 11.9 min. Figure S3 shows accompanying control experiments for the assays presented in Figure 2.

### Cryo-EM Structure Determination

Prior to grid preparation, PaFS_LL_ was dialyzed into cryo-EM buffer [50 mM HEPES (pH 7.5), 150 mM NaCl, 100 mM trimethylamine oxide (TMAO), and 1.5 mM TCEP] at 4°C for 1 h. MgCl_2_, geranylgeranyl thiodiphosphate (GGSPP), and NP-40 nonionic protein detergent were then added to final concentrations of 2.5 mM, 1 mM, and 0.001%, respectively. The final protein concentration was approximately 0.8 mg/mL. The sample was filtered by centrifugation in a 0.22-µM filtration unit (Millipore). R1.2/1.3 200-mesh copper grids (Quantifoil) were glow-discharged for 2 min (easiGlow, Pelco) prior to application of 3 µL protein sample. Grids were blotted for 4.5 s with a blot force of 0 at 100% humidity and flash-frozen in liquid ethane using a Mark IV Vitrobot (ThermoFisher Scientific). Frozen grids were clipped and transferred to a Titan Krios G3i cryogenic transmission electron microscope (Thermo Fischer Scientific) operating at 300 keV. Images were recorded with a K3 Summit electron detector at 81,000 magnification (0.54 Å/pixel) with a defocus of -1.0 to -3.0 µM. 40 frames were taken at a dose rate of 3.2-3.7 e^-^ /pixel/s (43 e^-^/Å^-2^). A total of 5,724 movies were collected from two identically prepared grids.

The PaFS_LL_ dataset was processed using cryoSPARC (version 4.4.1).^48^ After motion andCTF correction, 166 exposures were rejected during curation to yield 5,558 micrographs. Topaz Extract (box size of 384 Å) was used to generate a pool of 408,639 particles that were subjected to 2D classification (150 classes). Selection of all protein-resembling classes yielded a pool of 299,244 particles that were used to generate an *ab initio* 3D reconstruction into 5 different models. After two rounds of heterogenous refinement, the best model was used with 1,000 of the micrographs to train a Topaz model. The model was then used for re-extraction of the entire dataset, yielding a pool of 323,450 particles. 2D classification of these particles allowed for selection of 270,039 particles for further processing.

With these particles, we generated *ab initio* 3D reconstructions of eight different classes and subjected each class to four rounds of heterogenous refinement. After the refinement, four of the models, comprising 18,632 particles, contained only junk particles. One model (30,712 particles) clearly depicted the hexamer, with all six cyclase domains visible (one flat cyclase dimer and two tilted cyclase dimers). The remaining three classes, together containing 220,223 particles, consisted of octamer particles exclusively. Non-uniform refinement with *C*2 symmetry aligned these particles to a resolution of 3.56 Å. Particle subtraction of pendant cyclase domains and local refinement yielded a map with a resolution of 3.53 Å. DeepEMhancer was used to polish the final map of the octameric prenyltransferase core.^49^

The final model of the PaFS_LL_ prenyltransferase octamer (PDB 9B3T) contains 8 polypeptide chains and no ligands or water molecules. Chain A contains residues 374-669. Chain B contains residues 371-671. Chain C contains residues 374-672. Chain D contains residues 374-671. Chain E contains residues 374-672. Chain F contains residues 371-671. Chain G contains residues 374-672. Chain H contains residues 374-671. Several disordered loops were not modeled due to insufficient cryo-EM density. In chain A, the unmodeled residues are 407-409, 532-543, 565-581, 593-595, and 611-614. In chain B, the unmodeled residues are 531-543, 564-585, 593-596, and 610-614. In chain C, the unmodeled residues are 407-408, 531-543, 564-581, 610-615, and 655-657. In chain D, the unmodeled residues are 407-408, 531-542, 564-585, 593-596, 610-614, and 655-656. In chain E, the unmodeled residues are 407-410, 532-543, 565-581, 593-596, 611-614, and 653-657. In chain F, the unmodeled residues are 531-542, 564-583, 593-596, and 610-614. In chain G, the unmodeled residues are 407-408, 531-543, 566-581, 610-611, and 655-657. In chain H, the unmodeled residues are 407-408, 531-542, 564-585, 592-596, and 609-614. Additionally, residues 530-545 form a disordered loop connecting two helices; this loop was left unmodeled despite the presence of a patch of density disconnected from the main body of the structure in all eight chains.

The same pool of octamer particles was then subjected to 3D classification without a focus mask into 3–8 classes. However, this approach failed to produce a class with all eight cyclases in the flat orientation, so we subjected the same pool of particles to four rounds of heterogenous refinement with either 10, 12, 16, or 20 classes. A class with all eight cyclases in the flat orientation did not emerge. We then turned to 3D variability analysis with the pool of 220,223 octameric particles and generated a series of 20 maps that successfully accounted for the entire set of particles. We used UCSF ChimeraX to generate movie (.mov) files showing the transition across the set of 20 maps from different orientations, and converted the .mov files to GIFs using a freely available online tool (ezgif.com).

The 3D classification of the 220,223 octamer particle set with a focus mask over a cyclase pair dramatically improved the resolution of the cyclase domains. However, alignment of one particular cyclase pair for each octamer led to a decrease in clarity of the other three cyclase pairs, since many cyclase orientations were averaged together in those positions. Five classes were generated – the three disordered cyclase pairs were removed from each class using particle subtraction and the map was subjected to local refinement. Three of the five classes did not show clear density for the cyclase pair, while one class clearly showed a tilted cyclase pair, although the density was not strong enough to define an unambiguous orientation for the cyclase domain. One class (47,363 particles) revealed a flat cyclase pair at an overall resolution of 4.79 Å, with sufficiently well-defined cyclase densities to satisfactorily dock the crystal structure of the cyclase (PDB 5ERM^21^). The workflow of 3D reconstructions is summarized in Figure S7.

### PaFS_LL_ Model Building and Refinement

Atomic coordinates from the octameric prenyltransferase core of PaFS (PDB 8EAX) were docked into the 3D reconstruction of the PaFS_LL_ octameric prenyltransferase core using UCSF ChimeraX 1.3.^50^ Iterative rounds of refinement and model building were then performed with Phenix Real Space Refine^51^ and WinCoot.^52^ For real-space refinement, the resolution was set to 3.53 Å. Model quality was assessed with MolProbity.^53^

To build the model containing the ordered cyclase pair, the model of the octameric core refined against the PaFS_LL_ map was docked into the 3D reconstruction using UCSF ChimeraX 1.3. Two copies of the PaFS cyclase crystal structure (PDB 5ERM) were docked into the density for the cyclase pair also using UCSF ChimeraX 1.3. The ten chains were then opened alongside the map in WinCoot, and the linker region (PTQ) was added to connect the C-terminus of each cyclase domain to the N-terminus of the appropriate prenyltransferase domain. The resulting model, containing 8 chains, was used to create Figure 5.

### Mass Photometry and Sample Crosslinking

Each enzyme was diluted to approximately 50 nM (0.0008 mg/mL) in MP buffer [10 mM HEPES, pH 7.5, and 50 mM NaCl]. The mass photometer was calibrated using a standard curve derived from measurements of β-amylase and thyroglobulin, which allowed for the conversion of ratiometric contrast to mass in kDa. In a typical measurement, 12 µL of MP buffer were added to the drop receptacle to achieve the optimal focus position, and then 3 µL of protein was added into the drop to give a final protein concentration of approximately 10 nM. Data were collected for 60 s (approximately 60,000 frames) and analyzed using Refeyn AcquireMP software. For the crosslinking experiment, 0.2 % glutaraldehyde was added to PaFS_LL_ at 1.4 μM and allowed to react on ice for 10 min. The reaction was then quenched by addition of Tris buffer (pH 8.0) to a final concertation of 100 mM.

---

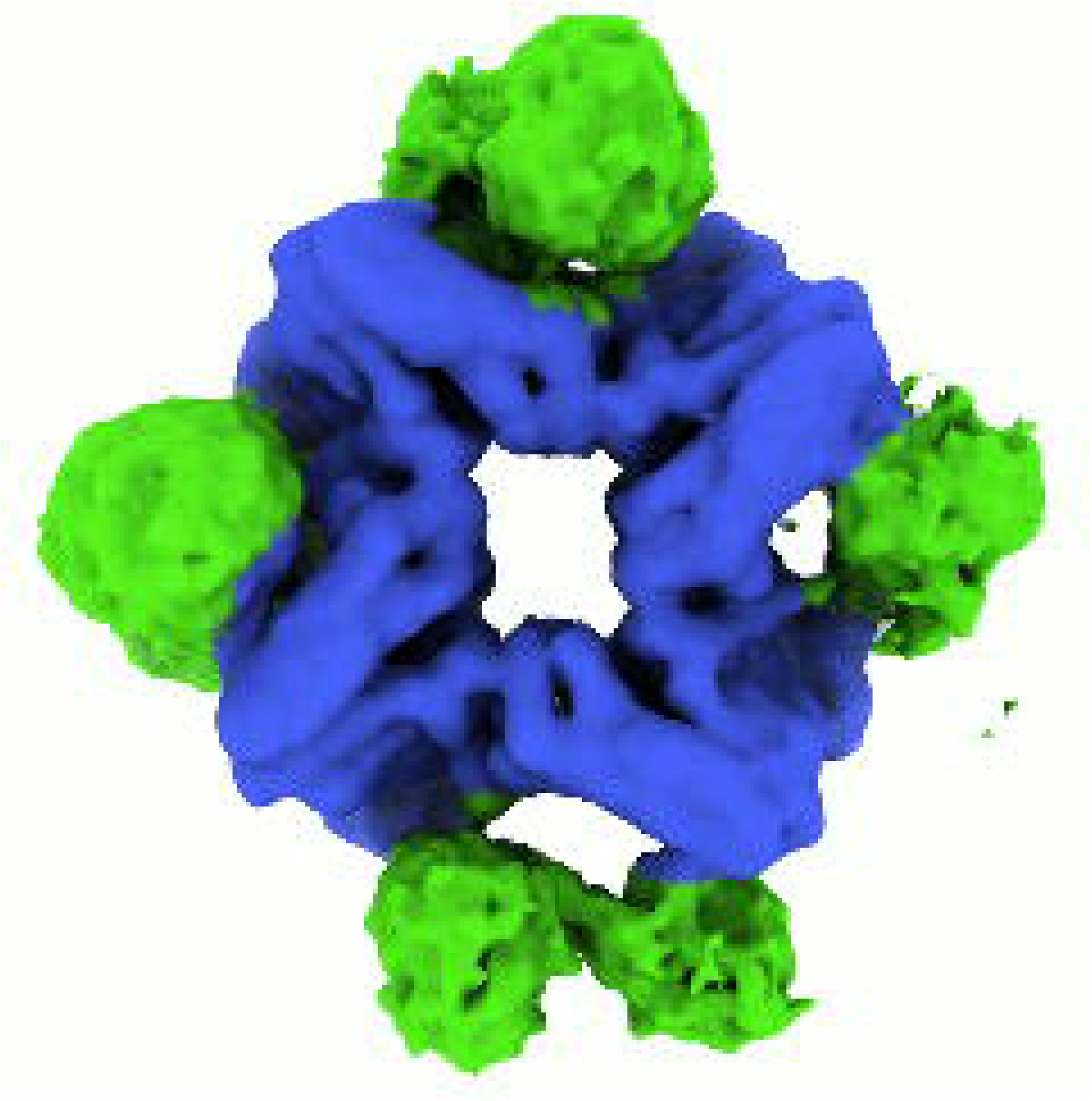

