## Supporting Information for "Engineering Substrate Channeling in Assembly-Line Terpene Biosynthesis"

### Table S1. Catalytic Parameters of Diterpene Cyclases

| **Catalytic Parameter** | **PaFS** | **$PaFS_{CY}$** | **$PaFS_{LL}$** | **SpS** | **CotB2** | **CotB2-$PaFS_{PT}$** |
|---|---|---|---|---|---|---|
| $V_{max}$ (μM $s^{-1}$) | 0.047 | 0.040 | 0.041 | 0.033 | 0.047 | 0.045 |
| $k_{cat}$ ($s^{-1}$) | 0.093 | 0.079 | 0.081 | 0.067 | 0.093 | 0.090 |
| $K_M$ (μM) | 8.60 | 6.12 | 8.03 | 6.74 | 12.26 | 9.24 |
| $k_{cat}/K_M$ ($M^{-1}s^{-1}$) | 10800 | 13000 | 10100 | 9890 | 7560 | 9730 |
| $R^2$ | 0.89 | 0.81 | 0.88 | 0.88 | 0.93 | 0.90 |
| Vmax (95% confidence interval) | 0.041-0.054 | 0.035-0.046 | 0.036-0.047 | 0.030-0.038 | 0.041-0.053 | 0.040-0.052 |
| $K_M$ (95% confidence interval) | 5.56-13.08 | 3.56-10.09 | 5.13-12.38 | 4.39-10.10 | 8.61-17.38 | 6.04-13.95 |

**Table S2. Cryo-EM data collection, reconstruction, and refinement statistics**

| | **$PaFS_{LL}$ Octameric Core** | **$PaFS_{LL}$ Core with Two Associated Cyclase Domains** |
|---|---|---|
| Magnification | 81,000 | 81,000 |
| Voltage (kV) | 300 | 300 |
| Exposure ($e^-$/Å$^2$) | 43 | 43 |
| Defocus range (µM) | -1.0 to -3.0 | -1.0 to -3.0 |
| Pixel size (Å/pix) | 0.54 | 0.54 |
| Symmetry imposed | C2 | C1 |
| Initial particles (no.) | 408,639 | 408,639 |
| Final particles (no.) | 220,663 | 47,363 |
| Map resolution (FSC=0.143) (Å) | 3.53 | 4.79 |
| Model | | |
| Initial model used | PDB 8EAX | |
| Model composition (#) | | |
| Chains | 8 | |
| Atoms | 16389 | |
| Residues | 2049 | |
| Water | 0 | |
| Ligands | 0 | |
| Root-mean-squared deviations | | |
| Bond lengths (Å) | 0.003 (0) | |
| Bond angles (°) | 0.615 (17) | |
| Validation | | |
| MolProbity score | 2.14 | |
| Clash score | 11.95 | |
| Poor rotamers (%) | 2.81 | |
| Ramachandran plot (%, MolProbity) | | |
| Favored | 96.67 | |
| Allowed | 3.33 | |
| Outliers | 0.00 | |
| Peptide plane (%) | | |
| Cis proline/general | 0.0/0.0 | |
| Twisted proline/general | 0.0/0.0 | |
| B-factors (Å$^2$) | | |
| (min/max/mean) | 30/113/65 | |
| Model vs. Data | | |
| CC (mask) | 0.73 | |
| CC (box) | 0.74 | |
| CC (peaks) | 0.72 | |
| CC (volume) | 0.73 | |
| PDB accession code | 9B3T | |
| EMDB accession code | EMD-44155 | EMD-44098 |

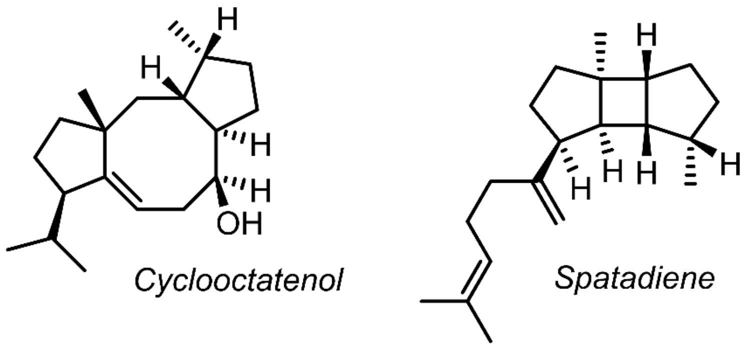


**Figure S1.** Structures of the products formed by the Class I diterpene cyclases CotB2 (left) and SpS (right).

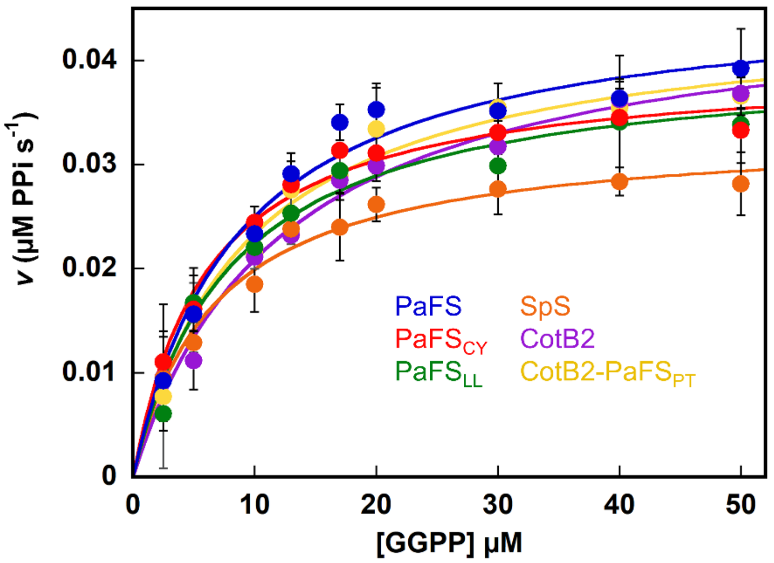


**Figure S2. Steady-state kinetics of diterpene cyclases.** Each datapoint is the initial velocity in a reaction with the specified enzyme and GGPP concentration and is the average of two identical trials. The curves were fit with the Michaelis-Menten equation to yield the catalytic parameters shown in Table S1. In every experiment, the enzyme concentration was 500 nM and the temperature was 25 °C.

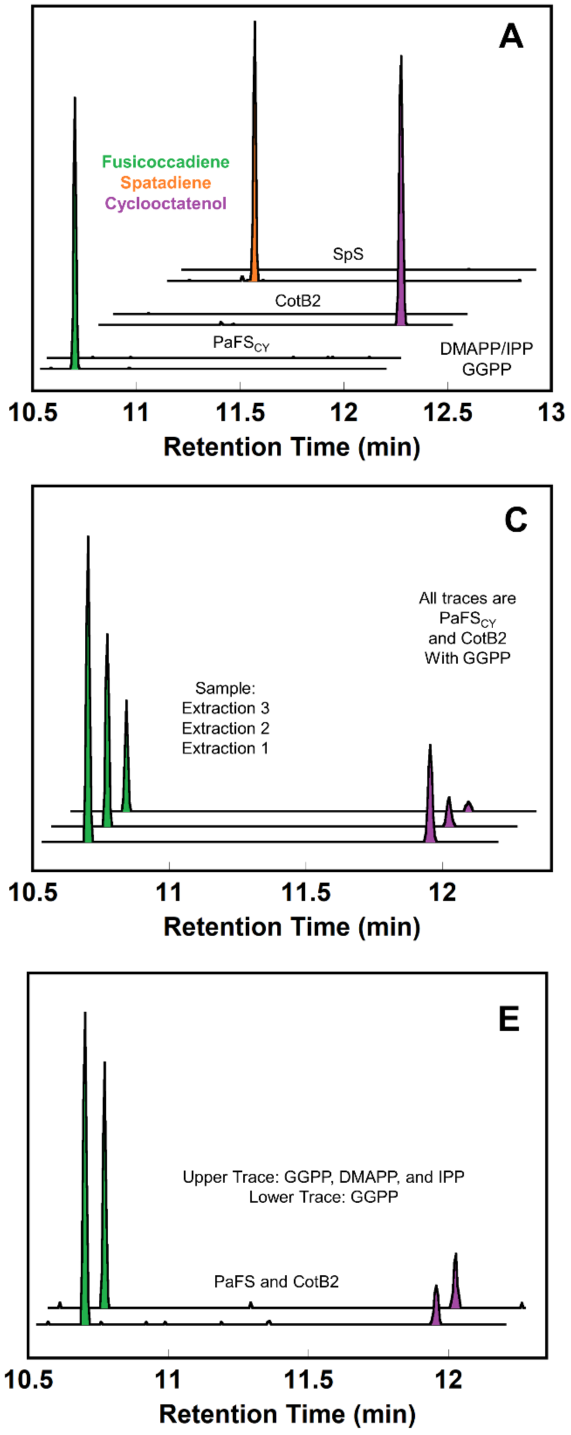


**Figure S3. Control experiments verifying that all enzyme constructs utilized produce the expected products from the expected substrates.** No other phenomenon is responsible for the observed fusicoccadiene enrichment attributed to substrate channeling between the prenyltransferase and cyclase domains of PaFS. (A) Each cyclase domain makes the expected product from GGPP and makes nothing from DMAPP and IPP. For each set of traces, the lower trace shows products formed from GGPP and the upper trace shows products formed from DMAPP and IPP. (B) Each bifunctional construct makes the expected product from GGPP or DMAPP and IPP, except that PaFS D474A makes nothing from DMAPP and IPP because its prenyltransferase domain is inactivated by the D474A substitution. For each set of traces, the lower trace shows products formed from GGPP and the upper trace shows products formed from DMAPP and IPP. (C) The ratio of products formed from GGPP does not change if the aqueous layer is extracted multiple times. (D) The ratio of products formed from GGPP does not change significantly over time. (E) IPP and DMAPP do not inhibit the cyclase domains differently (or at all). (F) The same ratio of products formed both from GGPP and from DMAPP and IPP is observed when the aqueous layer is extracted with different organic phases of differing polarities.

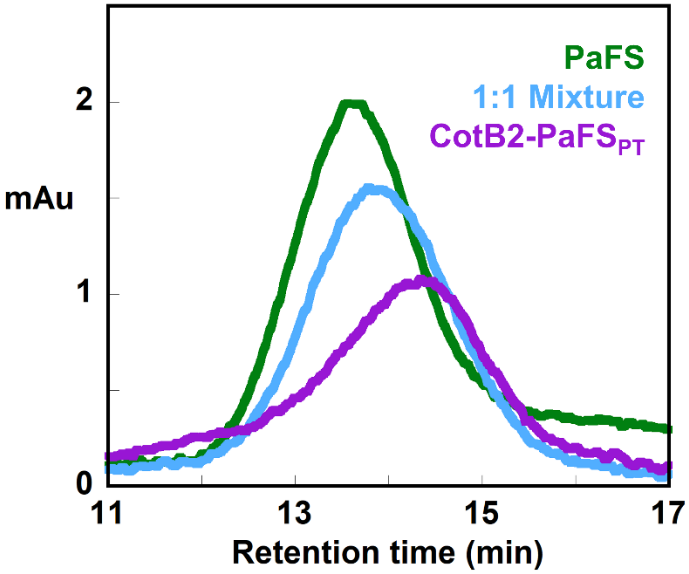


**Figure S4. Size-exclusion chromatography (SEC) of PaFS, CotB2-PaFS$_{PT}$, and an equimolar mixture of PaFS and CotB2-PaFS$_{PT}$.** The green chromatogram corresponds to PaFS when a 1 μM sample was applied to a 22-mL S600 column with a retention time of 13.6 min, roughly corresponding to the mass of an octamer (640 kD) based on a calibration curve for the column. The purple trace corresponds to a 1 μM sample of CotB2-PaFS$_{PT}$, which yields a retention time of 14.3 min. The CotB2-PaFS$_{PT}$ octamer is 32 kD smaller than the wild-type PaFS octamer, consistent with the slightly longer retention time. The light blue trace corresponds to a 1 μM sample of an equimolar mixture of PaFS and CotB2-PaFS$_{PT}$, which yields a retention time of 13.9 min. The intermediate retention time reflects the equilibration of the two proteins to form heterooctamers. No other peaks are visible on the chromatogram; suggesting that the protein concentration or some other aspect of the micro-environment of the SEC column stabilizes the higher-order oligomers.

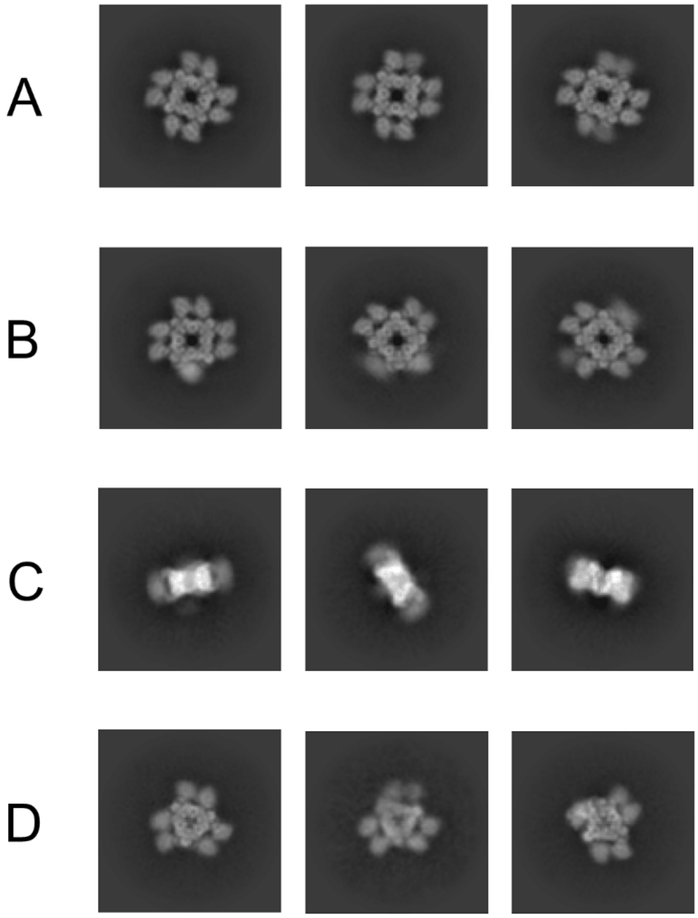


**Figure S5. Representative 2D classes of PaFS$_{LL}$.** (A) Top-down views of octamers in which cyclase pairs are approximately coplanar with each central prenyltransferase octamer. Some classes reveal a slight shading of one cyclase relative to the other in a given cyclase pair, suggesting that the cyclase pair is slightly tilted. (B) Top-down view of octamers in which some cyclase pairs are tilted. The tilting of cyclase pairs does not appear to be correlated – particles are observed with all four cyclase dimers flat (A), one tilted and three flat (B), and two tilted and two flat (*cis* or *trans*) (B). (C) Side view of octamers. (D)Top-down views of hexamers in which some cyclase pairs are tilted. The octamer:hexamer ratio is 220,000:31,000. If PaFS$_{LL}$ octamers and hexamers are in equilibrium, then the octamer:hexamer ratio would correspond to a $\Delta G$ of only 1.2 kcal/mol.

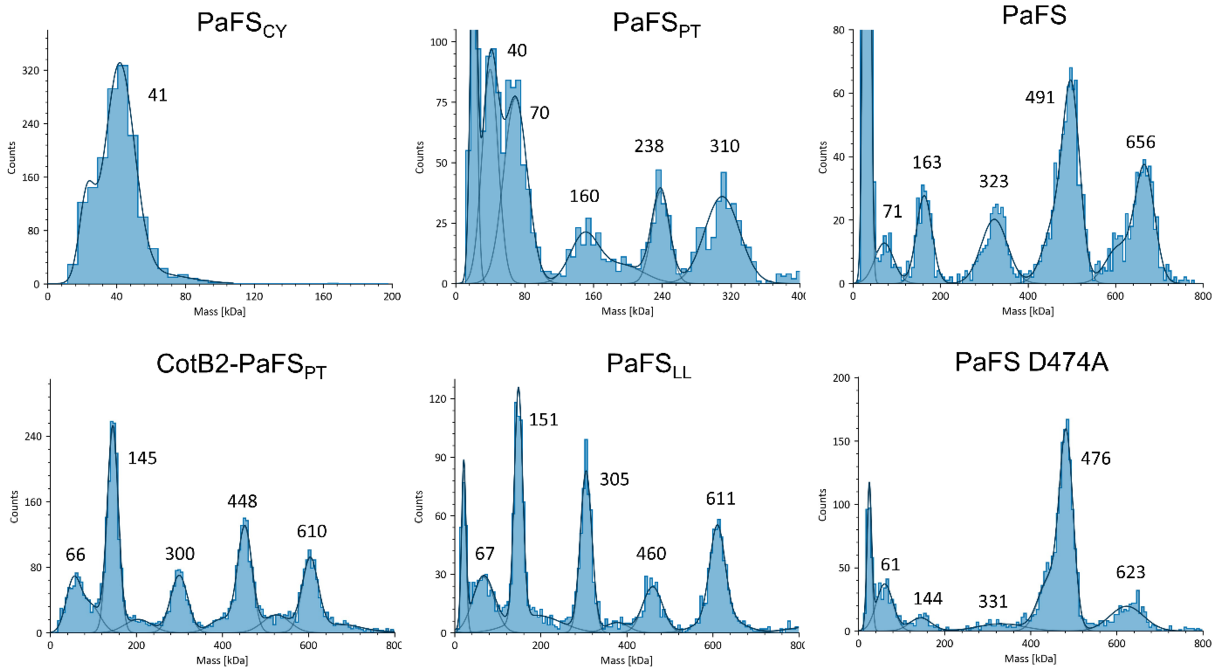


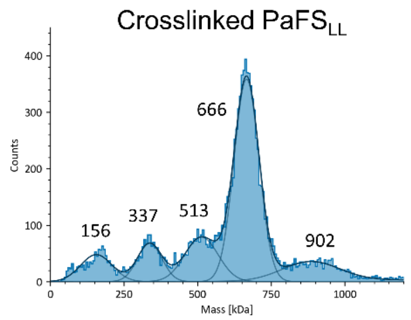


| Protein | Actual Mass (kDa) | x2 | x4 | x6 | x8 |
|---|---|---|---|---|---|
| PaFS$_{PT}$ | 38 | 76 | 152 | 228 | 304 |
| PaFS | 82 | 164 | 328 | 492 | 656 |
| CotB2-PaFS$_{PT}$ | 77 | 154 | 308 | 461 | 616 |
| PaFS$_{LL}$ | 75 | 150 | 300 | 450 | 600 |

**Figure S6. Oligomeric states of PaFS constructs determined by mass photometry.** At the experimental limit of 10 nM protein concentrations (0.0008 mg/mL), these samples are at a $10^3$-fold lower concentration than samples used for cryo-EM. PaFS$_{CY}$ does not oligomerize, whereas all other constructs (which contain the PaFS$_{PT}$ domain) reveal a mixture of oligomeric states containing monomer, dimer, tetramer, hexamer, and octamer. Oligomeric heterogeneity (not observed in Figure S2) may arise in part from the extremely low protein concentration required for the mass photometry experiment. To confirm this assessment, we crosslinked PaFS$_{LL}$ with glutaraldehyde at a concentration relevant for cryo-EM and then diluted the sample for mass photometry – the sample showed a significant encrichment of octamer (lower left). In each plot, the y-axis represents the number of individual species observed at the molecular mass indicated on the x-axis. Peaks are labeled with their average masses (determined from Gaussian functions fit to the data), which can be compared with the actual mass of each oligomeric state for each protein listed in the table.

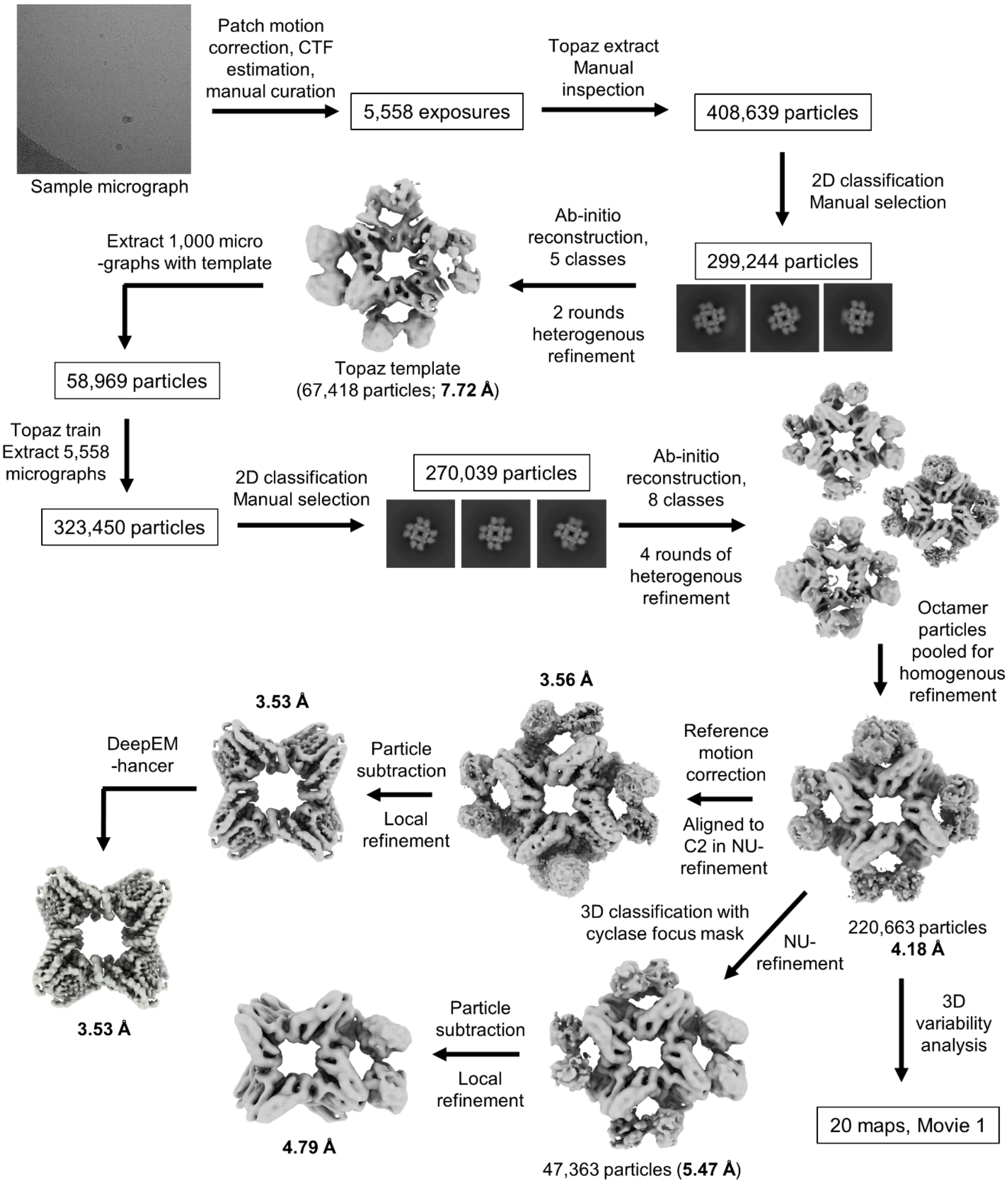


**Figure S7. Cryo-EM workflow.** A total of 5,558 exposures (sample micrograph shown) were taken from two grids. All data processing and analysis steps were conducted in cryoSPARC. Topaz was used to improve particle picking. Map sharpening was performed with DeepEMhancer.

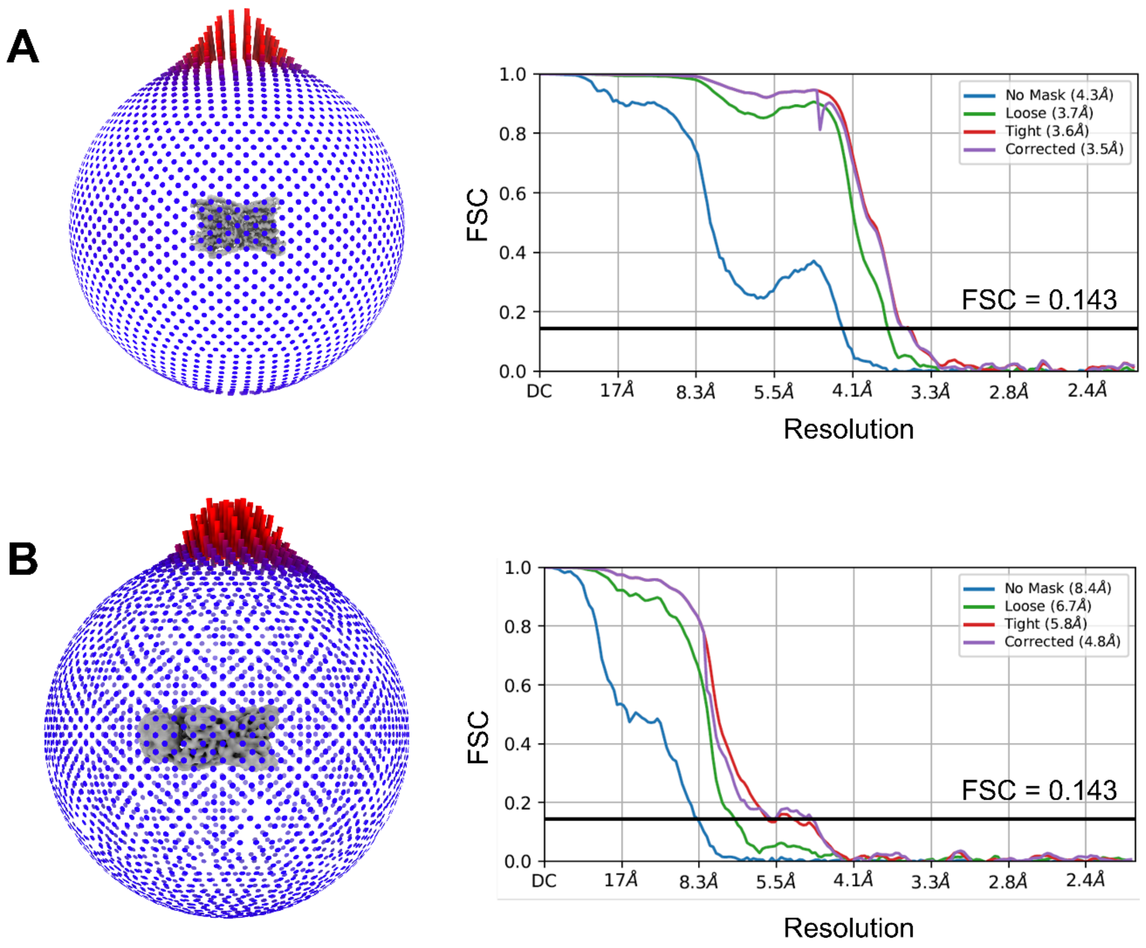


**Figure S8. FSC curves and data coverage plots for the Cryo-EM structures.** Angular distribution plots indicate a preferred "top-down" orientation of octamer particles in the dataset. (A) Angular distribution plot and Fourier shell correlation (FSC) plot for the structure of the octameric prenyltransferase core of $PaFS_{LL}$ with *C*2 symmetry. The resolution estimate at FSC = 0.143 is 3.53 Å. (B) Angular distribution and FSC plot for the map of the prenyltransferase core of $PaFS_{LL}$ with two associated cyclase domains. The resolution estimate at FSC = 0.143 is 4.79 Å.

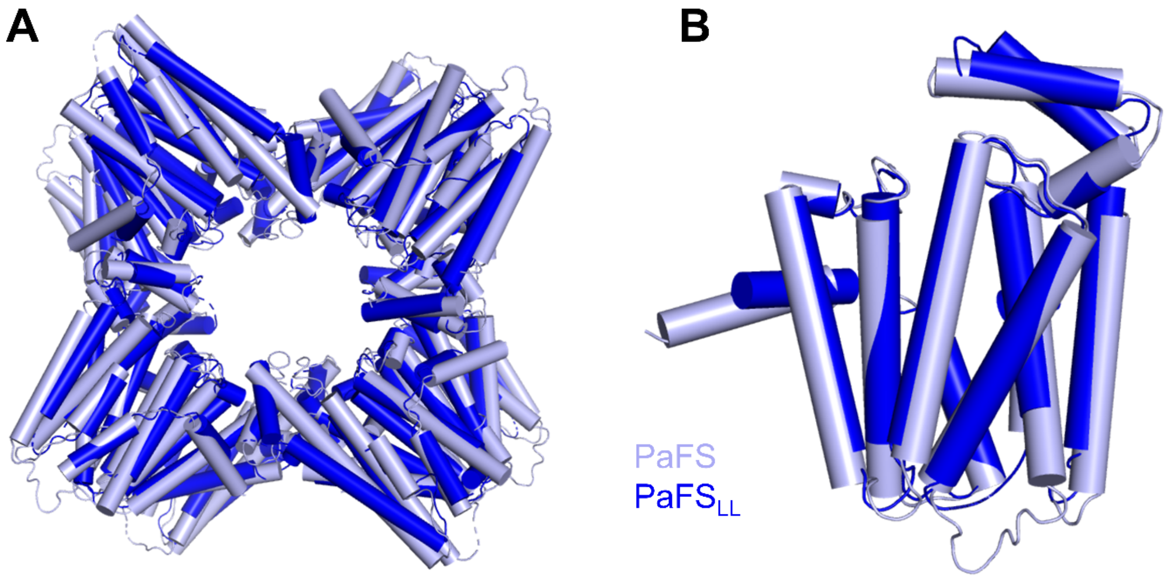


**Figure S9. Comparison of PaFS and PaFS$_{LL}$ prenyltransferase structures.** (A) Overlay of the PaFS (light blue; PDB 8EAX) and PaFS$_{LL}$ (dark blue) octameric prenyltransferase cores. (B) Overlay of chain A of PaFS and chain A of PaFS$_{LL}$, which aligned the most closely out of the eight chains.
